# The *SAMM50* rs3761472 causes mitochondrial dysfunction and metabolic dysfunction-associated steatotic liver disease

**DOI:** 10.1101/2025.06.04.657815

**Authors:** Suyeon Kim, Nahyun Kim, Uijin Kim, Young Jin Kim, Jun Ho Yun, Jiwon Heo, Hyunwoo Lee, Jiwon Choi, Inhae Jeong, Bong-Jo Kim, Ha Youn Shin

**Affiliations:** Department of Biomedical Science and Engineering, Konkuk University, Seoul, 05029, Republic of Korea; Terasaki Institute for Biomedical Innovation, Woodland Hills, 91367, California, United States of America; Division of Genome Science, Department of Precision Medicine, National Institute of Health, Cheongju-si, 28159, Republic of Korea; School of Advanced Biotechnology, Konkuk University, Seoul, 05029, Republic of Korea

**Keywords:** CRISPR/Cas9, Metabolic dysfunction-associated steatotic liver disease (MASLD), Precision medicine, SAMM50, Single nucleotide polymorphism (SNP)

## Abstract

Genome-wide association studies (GWAS) have identified the *SAMM50* rs3761472 single nucleotide polymorphism (SNP) as a risk factor for metabolic dysfunction-associated steatotic liver disease (MASLD), although its in vivo functions remain unclear. *SAMM50* encodes a mitochondrial outer membrane protein critical for maintaining mitochondrial structure. To investigate the biological effects of rs3761472, we generated *Samm50*-knock-in (KI) mice harboring a D110G substitution using CRISPR/Cas9. This variant impaired mitochondrial integrity by downregulating key regulators of mitochondrial architecture, dynamics, and quality control. This contributed to reduced ATP production and elevated oxidative stress, inflammation, and hepatocyte death. The mutation also induced insulin resistance and glucose intolerance. *Samm50*-KI mice fed a high-fat diet exhibited pronounced hepatic lipid accumulation and liver injury, highlighting its pathogenic role in MASLD progression. Our findings demonstrate that *SAMM50* rs3761472 is a critical driver of mitochondrial dysfunction and MASLD susceptibility, supporting its potential as a therapeutic target and its relevance to precision medicine.

## Introduction

Metabolic dysfunction-associated steatotic liver disease (MASLD) is one of the most prevalent chronic liver diseases worldwide and a growing global health concern. MASLD encompasses a spectrum of liver disorders ranging from simple steatosis to steatohepatitis, fibrosis, and cirrhosis, and may ultimately progress to hepatocellular carcinoma (Anstee & Day, 2013). MASLD development is influenced by environmental and genetic factors, with genetic variants playing a key role in determining individual susceptibility (Anstee & Day, 2013, Kim & Park, 2020). Despite the increasing prevalence of MASLD, research on its genetic factors has primarily focused on clinical correlations, leaving the molecular mechanisms underlying pathogenesis largely unexplored.

Genome-wide association studies (GWAS) have identified several genetic loci associated with MASLD susceptibility, most notably *PNPLA3*, *TM6SF2*, and *SAMM50* (Chen, Lin et al., 2015, Chung, Lee et al., 2018, Kitamoto, Kitamoto et al., 2013, Li, Shen et al., 2021, Luo, Oldoni et al., 2022, Qiao, Yang et al., 2022, Zhao, Xu et al., 2023). Unlike PNPLA3 and TM6SF2, which primarily regulate lipid metabolism, SAMM50 maintains mitochondrial structure, suggesting that its genetic variant may contribute to MASLD pathogenesis through distinct mechanisms (Luo et al., 2022, Mazo, Malta et al., 2019, Zhao et al., 2023). *SAMM50* encodes a key component of the sorting and assembly machinery (SAM) complex in the outer mitochondrial membrane (OMM), which is involved in the assembly of β-barrel proteins and the maintenance of mitochondrial architecture (Ott, Dorsch et al., 2015). Among four MASLD-associated single-nucleotide polymorphisms (SNPs) in *SAMM50* identified in the GWAS discovery set, rs3761472 harbors an amino acid change that significantly increases the susceptibility to MASLD (Chen et al., 2015, Chung et al., 2018, Kitamoto et al., 2013, Li et al., 2021, Qiao et al., 2022, Zhao et al., 2023). The *SAMM50* rs3761472 has a minor-allele frequency (MAF) of 39.9% in East Asians and 15.5% in Europeans (gnomAD v4.1), comparable to *PNPLA3* rs738409 (MAFEAS=41.4%, MAFEUR=21.8%) but markedly higher than *TM6SF2* rs58542926 (MAFEAS=7.7%, MAFEUR=7.4%). By directly influencing mitochondrial structure and function, the mutant protein encoded by this *SAMM50* SNP could promote MASLD pathogenesis, offering new insights into mitochondrial contributions to the disease. This is consistent with the increasing recognition of the importance of mitochondria in MASLD, reflecting their broader involvement in various diseases (Nassir & Ibdah, 2014, Tian, Liu et al., 2022).

Mitochondria are essential organelles that regulate ATP production and play critical roles in cellular respiration, lipid metabolism, cell signaling, reactive oxygen species (ROS) production, and apoptosis (McBride, Neuspiel et al., 2006). Aberrant mitochondrial function is closely linked to a range of disorders, including MASLD (Nassir & Ibdah, 2014). Given the critical role of mitochondria in MASLD development, understanding how genetic variants such as *SAMM50* rs3761472 influence mitochondrial function is crucial. In this study, we investigated the specific mechanisms using a knock-in (KI) mouse model harboring the corresponding D110G missense mutation in the *Samm50* gene, generated via the CRISPR/Cas9 system. Our findings provide the first in vivo evidence that the resulting SAMM50 D110G exacerbates MASLD by impairing mitochondrial function. By elucidating these molecular pathways, this study aims to lay the groundwork for developing novel therapeutic strategies targeting mitochondrial dysfunction in MASLD. This study not only deepens our understanding of the genetic basis of MASLD but also paves the way for the development of precision therapies specifically designed for individuals carrying the *SAMM50* rs3761472 variant.

## Results

### Generation of CRISPR/Cas9–mediated KI mice carrying the *Samm50* rs3761472 genetic variant

The *SAMM50* rs3761472 SNP resulted in a single amino acid change from aspartic acid to glycine at position 110, which is highly conserved across multiple species (Fig. 1A, B). This specific *SAMM50* SNP is predominantly found in Asian patients with MASLD, including Chinese Han, Japanese, and Korean populations (Chen et al., 2015, Chung et al., 2018, Kitamoto et al., 2013). It has also been identified among 28 SNPs from approximately 7,000 Koreans genotyped by the Korea Biobank Array as significantly associated with liver metabolism (Appendix Fig. S1). When we expanded the association analysis of hepatometabolic traits in 151,912 Korean Biobank participants, each copy of the rs3761472-G allele was significantly associated (P < 5x10^-8^) with higher serum liver enzymes (alanine aminotransferase, ALT: β = 0.011, P = 1.34×10^-99^; aspartate aminotransferase, AST: β = 0.007, P = 1.17×10^-102^), after adjusting for age, age², sex, BMI, recruitment region, and the first four genetic principal components. Biobank Japan PheWeb data further reveal that rs3761472 is associated with increased risk of cirrhosis (OR = 1.15; P = 1.00×10^-5^) and hepatocellular carcinoma (OR = 1.18; P = 1.19×10^-8^) (Sakaue, Kanai et al., 2021).

**Figure 1.**
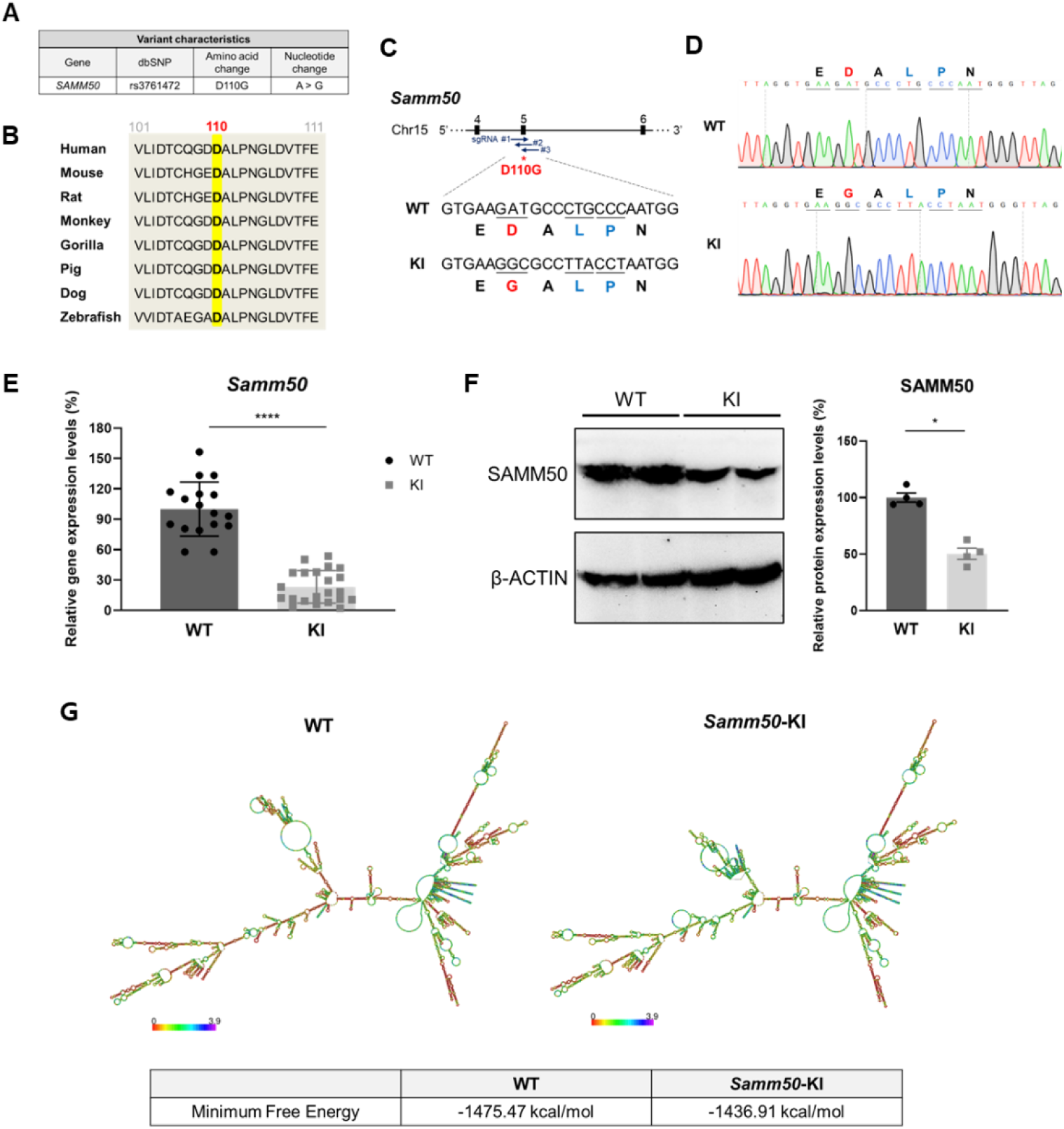
Characterization of *Samm50*-KI mice carrying rs3761472. (**A**) The major characteristics of the *SAMM50* rs3761472 SNP. (**B**) Alignment of amino acid sequences showing that aspartic acid (D) at position 110 in SAMM50 is highly conserved across various species. (**C**) Schematic illustration depicting the location of sgRNA binding to the target region and the amino acid sequence changes in *Samm50*-KI mice. In KI mice, D110 is converted to glycine (G) in exon 5 of the *Samm50* gene (indicated in red). Blue letters indicate a silent mutation introduced to prevent non-specific targeting by Cas9. This region contains NGG sequences, which can be non-specifically cut by the Cas9 endonuclease. (**D**) Sanger sequencing analysis confirming the homozygous mutation of rs3761472 in the *Samm50* gene of KI mice. (**E**) Relative *Samm50* gene expression levels in the livers of WT and *Samm50*-KI mice. mRNA levels of *Samm50* were normalized to *Gapdh* mRNA levels, and KI mRNA levels are presented relative to those of WT (WT, n = 17; *Samm50*-KI, n = 22; ****p < 0.0001). (**F**) SAMM50 protein expression levels in WT and *Samm50*-KI mice. GAPDH was used as a loading control (left panel). Quantification of SAMM50 protein expression levels (right panel) (WT, n = 4; *Samm50*-KI, n = 4). Results are presented as means ± SEM (*p < 0.05). (**G**) Comparison of centroid secondary structures of *Samm50* mRNA in WT and KI mice with minimum free energy. Data are presented as means ± SEM, and statistical significance was determined using the Mann–Whitney test (**E, F**).

To investigate the functional consequences of this genetic variant in vivo, we generated *Samm50*-KI mice harboring the D110G mutation using the CRISPR/Cas9 system (Fig. 1C; Fig. EV1A, B). To introduce the D110G mutation, we modified exon 5 of *Samm50* by substituting the original GAT codon with GGC. We also introduced silent mutations, CTG to TTA and CCC to CCT, in the adjacent target sequence to eliminate non-target protospacer adjacent motif (PAM) sequences, reducing the likelihood of off-target cleavage by Cas9 nuclease. Homozygous KI mice were screened and confirmed by PCR-based genotyping, followed by Sanger sequencing (Fig. 1D). Intriguingly, *Samm50* mRNA and SAMM50 protein levels were markedly reduced in *Samm50*-KI mouse livers, indicating the importance of D110 (Fig. 1E, F). To further explore how the D110G significantly affected expression levels, we predicted the secondary structure of *Samm50* mRNA (http://rna.tbi.univie.ac.at/) (Gruber, Lorenz et al., 2008). A comparative analysis revealed a distinct alteration in the centroid secondary structure of *Samm50*-KI mRNA relative to wild-type (WT) mRNA (Fig. 1G), which resulted in a 38.56 kcal/mol increase in minimum free energy, indicating reduced mRNA stability (Wu, Wang et al., 2022). This reduction in mRNA stability may explain the diminished *Samm50* expression in KI mice, further underscoring the functional importance of D110 residue.

### Impact of the SAMM50 D110G mutation on mitochondrial membrane components

Several lines of evidence indicate that *SAMM50* knockdown in vitro reduces the expression of mitochondria-associated components such as DNAJC11, MTX1, MTX2, OPA1, DRP1, and VDAC (Ioakeimidis, Ott et al., 2014, Kozjak-Pavlovic, Ross et al., 2007, Liu, Gao et al., 2016). To determine whether this effect manifests in vivo, we examined the impact of SAMM50 D110G on key mitochondrial genes maintaining structural integrity in *Samm50*-KI mice. Various protein complexes maintain the double-membrane structure of mitochondria, which comprises the OMM and inner mitochondrial membrane (IMM) (Fig. EV2A). These include translocase of the outer membrane (TOM) and SAM complexes, with SAMM50 as a core component. The mitochondrial contact site and cristae organizing system (MICOS) in the IMM forms contact sites that connect the OMM and IMM, and the mitochondrial intermembrane space bridging (MIB) complex encompasses both SAM and MICOS complexes (Huynen, Muhlmeister et al., 2016, Pfanner, Wiedemann et al., 2004). We found reductions in the expression of MIB components, including *Dnajc11*, *Mic19*, *Mic60*, *Mtx1*, and *Mtx2*, as well as the TOM members *Tom40* and *Tom70* in the livers of *Samm50*-KI (Fig. 2A, B).

**Figure 2.**
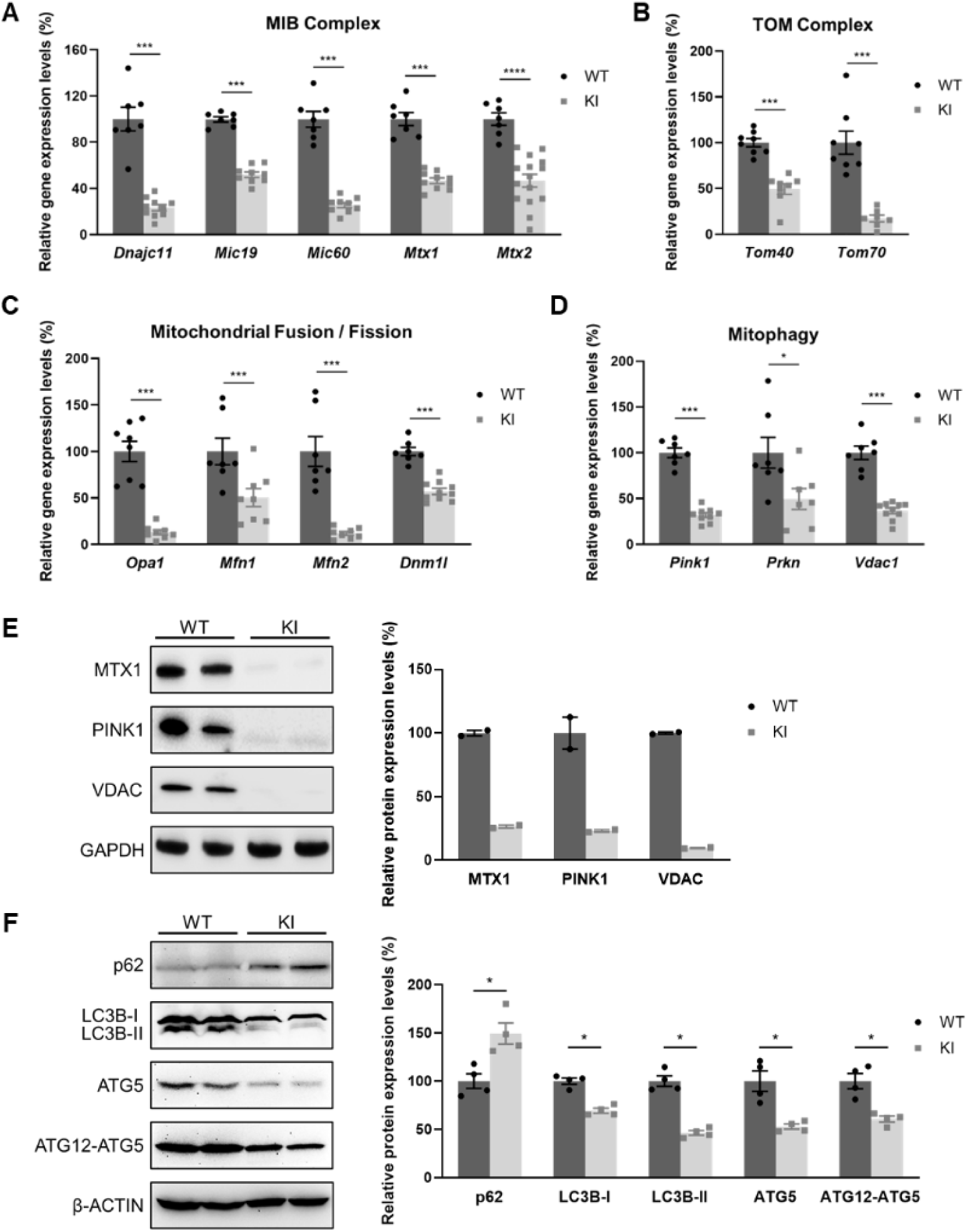
The impact of the *Samm50* genetic variant rs3761472 on mitochondrial membrane structure and functional integrity in mice. (**A, B**) Relative mRNA levels of genes associated with mitochondrial membrane protein complexes MIB (**A**) and TOM (**B**). (**C, D**) Relative mRNA levels of genes involved in mitochondrial fusion and fission (**C**) and mitophagy (**D**) (WT, n = 7; *Samm50*-KI, n ≥ 9 mice; ***p < 0.001; ****p < 0.0001). (**E**) Left: Western blot analyses of MTX1, PINK1, and VDAC protein expression levels. GAPDH was used as a loading control. Right: Quantification of MTX1, PINK1, and VDAC protein expression levels in WT and *Samm50*-KI mice. Relative protein expression levels in *Samm50*-KI mice were normalized to those in WT mice (WT, n = 2; *Samm50*-KI, n = 2). (**F**) Left: Western blot analyses of p62, LC3B, ATG5, and ATG12-ATG5 conjugate. GAPDH was used as a loading control. Right: Quantification of protein expression levels in WT and *Samm50*-KI mice. Relative protein expression levels in *Samm50*-KI mice were normalized to those in WT mice (WT, n = 4; *Samm50*-KI, n = 4; *p < 0.05). Data are presented as means ± SEM, and statistical significance was determined using the Mann– Whitney test (**A-D, F**).

Further in silico analyses revealed differences in the protein structures of the SAMM50 D110G mutant and SAMM50 WT (Fig. EV2B-D). Protein structures were predicted using ColabFold v1.5.5, which implements AlphaFold2 (Mirdita, Schutze et al., 2022). Structural comparison of SAMM50 WT and D110G showed notable differences, particularly in loop regions, suggesting that the D110G mutation may disrupt folding dynamics or structural stability. The RMSD, a standard metric for structural similarity, was 10.098 Å, indicating that conformational changes may compromise mitochondrial membrane integrity (Kosloff & Kolodny, 2008).

### SAMM50 D110G compromises mitochondrial dynamics and quality control

We further investigated whether the SAMM50 D110G variant affected genes involved in mitochondrial dynamics and quality control, as mitochondria are dynamic organelles that undergo fusion and fission to sustain their function (Youle & van der Bliek, 2012). OPA1 is crucial for fusion of the IMM, whereas mitofusins (MFN1 and MFN2) mediate fusion of the OMM (Song, Ghochani et al., 2009). DRP1 is a key regulator of mitochondrial fission (Ikeda, Shirakabe et al., 2015). We found that the mRNA levels of *Opa1*, *Mfn1*, *Mfn2*, and *Dnm1l* were significantly reduced in the livers of *Samm50*-KI mice (Fig. 2C).

Mitophagy is essential for mitochondrial quality control by removing damaged mitochondria. Under physiological conditions, PINK1 is imported into mitochondria and degraded, but upon mitochondrial damage, it accumulates on the outer membrane, recruits Parkin, and promotes ubiquitination of proteins such as VDAC (Springer & Kahle, 2011, Sun, Vashisht et al., 2012). Ubiquitinated proteins are recognized by p62 and delivered to autophagosomes via LC3B (Mizushima & Komatsu, 2011). Notably, certain forms of mitochondrial dysfunction can impair mitophagy, leading to the accumulation of defective mitochondria and increased cellular stress (Song, Franco et al., 2017). In *Samm50-*KI mouse livers, the mRNA levels of *Pink1*, *Prkn*, and *Vdac1*, genes critical for mitophagy initiation, were reduced, indicating impaired activation of the mitophagy pathway (Fig. 2D). Consistent with these observations, protein levels of PINK1, VDAC, and MTX1 were also decreased (Fig. 2E). To further assess whether mitophagic flux was impaired in *Samm50*-KI, we examined key autophagy-related markers, p62, LC3B, ATG5, and the ATG12–ATG5 conjugate. p62 serves as an autophagy adaptor and accumulates when autophagic degradation is impaired, while LC3B and ATG5 are essential for autophagosome formation. In *Samm50*-KI, p62 was increased, while LC3B, ATG5, and the ATG12–ATG5 conjugate were reduced (Fig. 2F). Collectively, these findings indicate that the SAMM50 D110G mutation disrupts mitochondrial dynamics and mitophagy, thereby compromising mitochondrial quality control in the liver.

### A single amino acid change, D110G, impairs the mitochondrial function of SAMM50

We next investigated the functional consequences of the SAMM50 D110G on mitochondrial processes, including calcium homeostasis, ATP production, and oxidative metabolism. Mitochondria regulate calcium signaling through uptake by the mitochondrial calcium uniporter (MCU) complex, thereby influencing ATP synthesis. In *Samm50*-KI, the expression of MCU complex–related genes was reduced (Fig. 3A). In addition, genes encoding components of the oxidative phosphorylation (OXPHOS) system—including electron transport chain Complexes I–IV and Complex V (ATP synthase)—were downregulated (Fig. 3B). Concomitantly, NAD^+^/NADH ratios were decreased (Fig. 3C, D), and hepatic ATP levels were markedly reduced by approximately 50% in *Samm50*-KI compared with WT controls (Fig. 3E).

**Figure 3.**
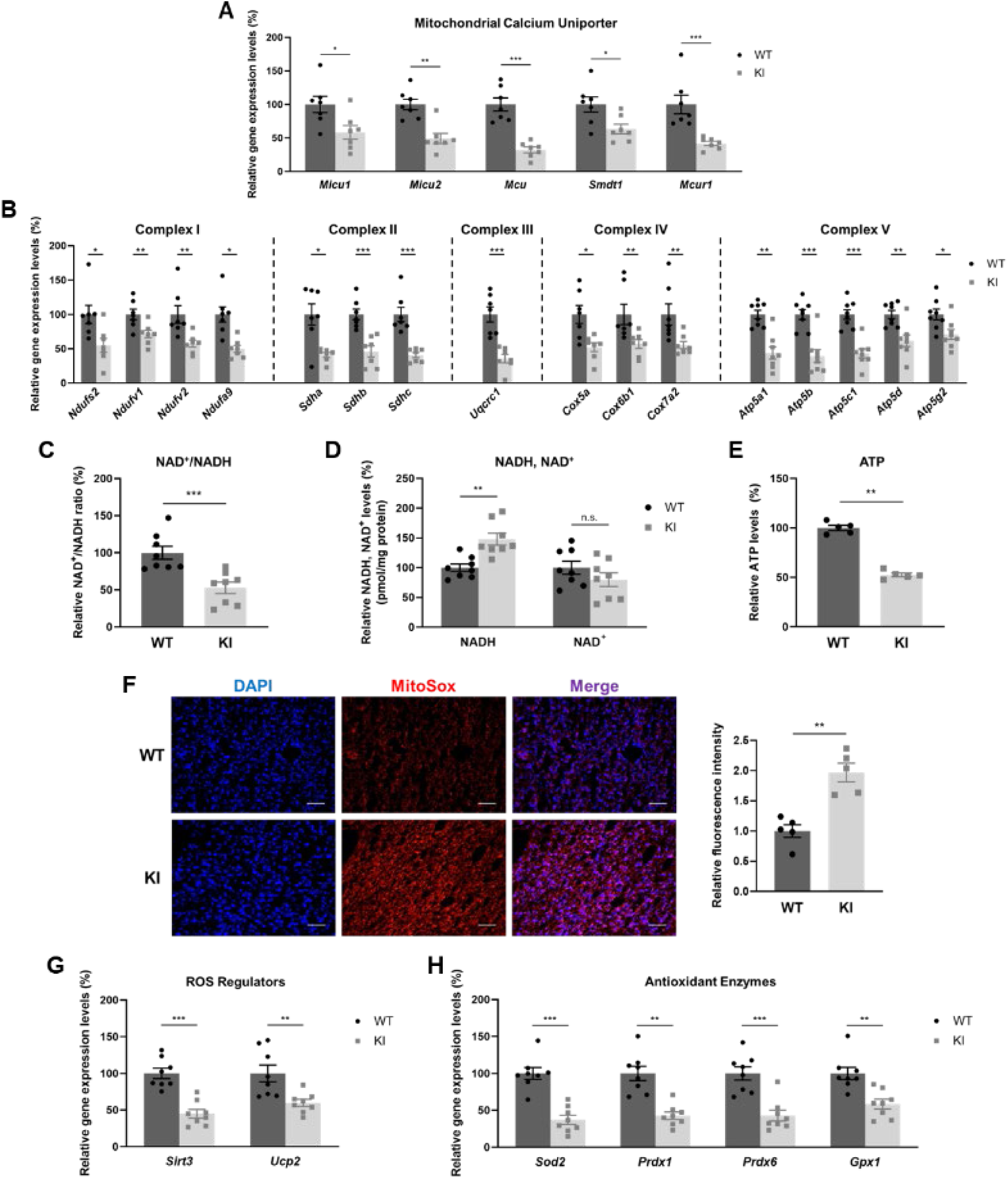
*Samm50*-KI mice display mitochondrial dysfunction. (**A, B**) Relative mRNA levels of genes associated with mitochondrial calcium uniporters (A) and OXPHOS complexes (**B**) (WT, n ≥ 7; *Samm50*-KI, n ≥ 7; *p < 0.05; **p < 0.01; ***p < 0.001). (**C, D**) Comparison of NAD^+^/NADH ratio (**C**) and NADH and NAD^+^ concentrations (**D**) in the livers of WT and *Samm50*-KI mice (WT, n = 8; *Samm50*-KI, n = 8; **p < 0.01; ***p < 0.001; n.s., not significant). (**E**) Hepatic ATP levels of WT and *Samm50*-KI mice (WT, n = 5; *Samm50*-KI, n = 5; **p < 0.01) (**F**) Left: Representative fluorescence images of mitochondrial ROS in frozen liver tissue, stained with MitoSOX Red (red). Cell nuclei were counterstained with DAPI (blue) and visualized using fluorescence microscopy (magnification, 200x). Scale bars: 100 μm. Right: Quantitative analysis of the relative fluorescence intensity of MitoSOX Red in WT and *Samm50*-KI mouse livers (WT, n = 5; *Samm50*-KI, n = 5; **p < 0.01). (**G, H**) Relative gene expression levels of ROS regulators (**G**) and antioxidant enzymes (**H**) (WT, n = 8; *Samm50*-KI, n = 8; **p < 0.01; ***p < 0.001). mRNA levels of each gene were normalized to those of *Gapdh*, and KI mRNA levels are presented relative to those of WT. Data are presented as means ± SEM, and statistical significance was determined using the Mann– Whitney test (**A-H**).

Mitochondrial dysfunction increases mitochondrial ROS (mtROS) levels owing to impaired electron transport and leakage, triggering mitochondrial damage and cell death (Ott, Gogvadze et al., 2007). Consistently, mtROS levels were increased in *Samm50*-KI (Fig. 3F; Appendix Fig. S2), accompanied by a significant reduction in the expression of antioxidant-related genes, including *Sirt3*, *Ucp2*, *Sod2*, *Prdx1*, *Prdx6*, and *Gpx1* (Fig. 3G, H). Taken together, SAMM50 D110G inhibits ATP production and elevates mtROS levels, ultimately compromising mitochondrial function.

### SAMM50 D110G mutation triggers inflammation and cell death in mouse liver

As mitochondrial dysfunction induces inflammation through oxidative stress and cytokine release, we evaluated the expression levels of immune-related genes. This analysis showed elevated expression of the NF-κB subunits *Nfkb2* and *RelB*, the pro-inflammatory cytokines *Il6*, *Tnfa*, and *Il1b,* and type I and II interferons *Ifna*, *Ifnb*, and *Infg* (Fig. 4A-C). In addition, elevated levels of *Nos2* and *Cox2* mRNA and NOS2 protein were observed in *Samm50*-KI (Fig. 4D, E).

**Figure 4.**
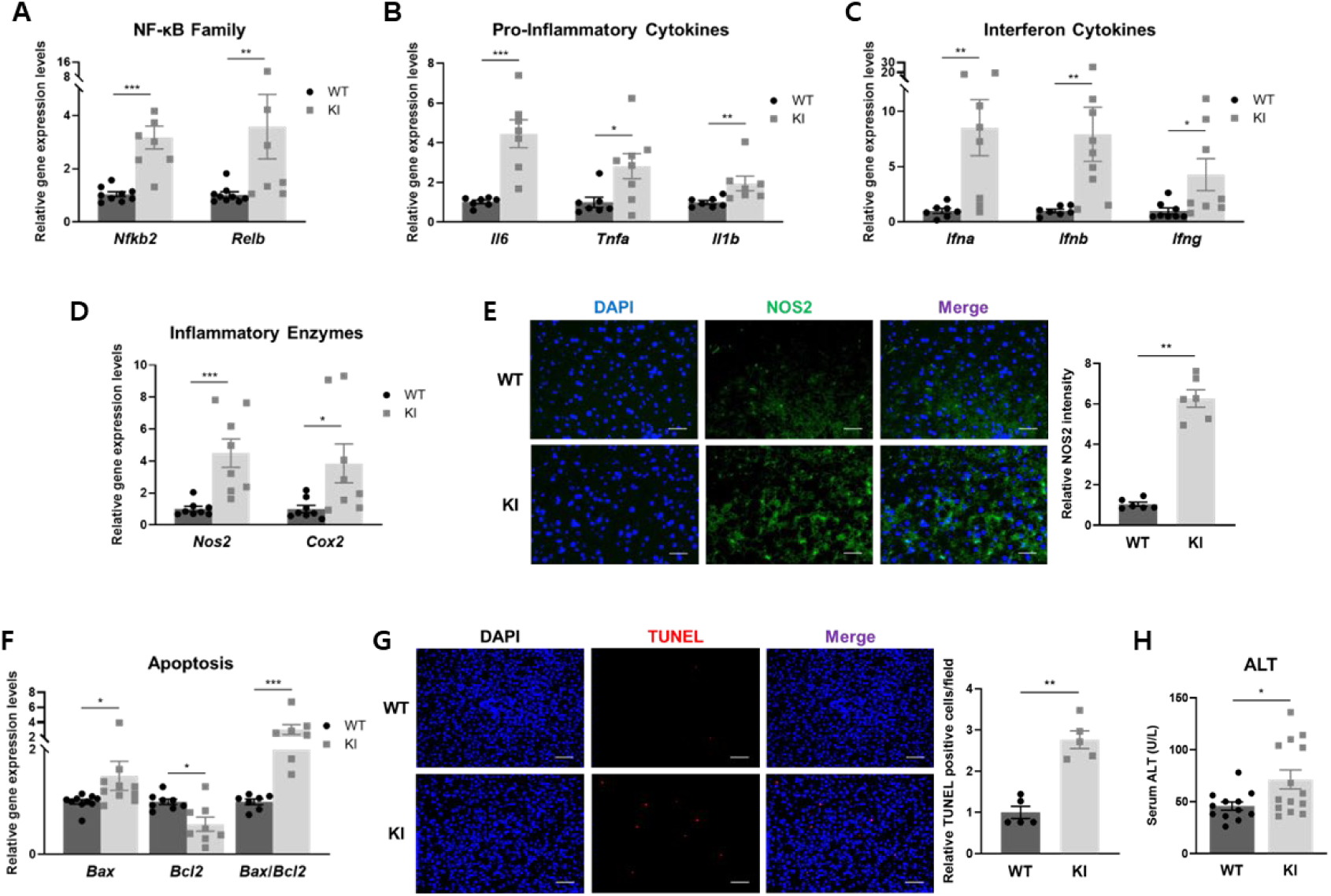
SAMM50 D110G induces cytokine gene expression and hepatocyte death. (**A-D**) Relative gene expression levels of NF-κB subunits (**A**) (WT, n = 7; *Samm50*-KI, n = 8; **p < 0.01; ***p < 0.001), pro-inflammatory cytokines (**B**) (WT, n = 7; *Samm50*-KI, n = 7; *p < 0.05; **p < 0.01; ***p < 0.001), type 1 and type 2 interferons (**C**) (WT, n ≥ 7; *Samm50*-KI, n ≥ 7; *p < 0.05; **p < 0.01), and *Nos2* and *Cox2* (**D**) (WT, n = 8; *Samm50*-KI, n = 8; *p < 0.05; ***p < 0.001). (**E**) Left: Immunofluorescence analysis of NOS2 proteins in the livers of WT and *Samm50*-KI mice (magnification, 400x). Scale bars: 100 μm. Right: Quantitative analysis of the relative fluorescence intensity of NOS2 proteins (WT, n = 6; *Samm50*-KI, n = 6; **p < 0.01). (**F**) Relative gene expression levels of *Bax* and *Bcl-2* and the relative *Bax*/*Bcl-2* ratio (WT, n ≥ 7; *Samm50*-KI, n ≥ 7; *p < 0.05; ***p < 0.001). (**G**) Left: Detection of in situ DNA fragmentation in cell nuclei by the TUNEL assay (red). DAPI (blue) and visualized using fluorescence microscopy (magnification, 200x). Scale bars: 100 μm. Right: Quantification of the relative number of TUNEL-positive cells per field (WT, n = 5; *Samm50*-KI, n = 5; **p < 0.01). (**H**) Serum ALT levels of WT and *Samm50*-KI mice (WT, n = 12; *Samm50*-KI, n = 14; *p < 0.05). mRNA levels of each gene were normalized to those of *Gapdh*, and KI mRNA levels are presented relative to those of WT. Data are presented as means ± SEM, and statistical significance was determined using the Mann–Whitney test (**A-H**).

Mitochondrial dysfunction and inflammation can accelerate cell death. We measured the expression of *Bax* and *Bcl2*, which regulate mitochondrial membrane integrity. Bax promotes apoptosis by increasing membrane permeability, whereas Bcl2 opposes this effect and supports cell survival (Hsu & Youle, 1997). In *Samm50*-KI livers, mRNA levels of *Bax* were increased and *Bcl2* reduced, resulting in an elevated *Bax*/*Bcl2* ratio (Fig. 4F), along with a higher number of TUNEL-positive cells (Fig. 4G). Consistent with these findings, ALT levels were significantly elevated in *Samm50*-KI, indicating liver damage (Fig. 4H). Additionally, AST levels showed a ∼30% increase (Appendix Fig. S3). Taken together, mitochondrial dysfunction caused by the SAMM50 D110G mutation triggers an immune response, ultimately leading to cell death and liver damage.

### SAMM50 D110G causes moderate insulin resistance and glucose intolerance

Mitochondrial dysfunction is also known to precede insulin resistance in MASLD (Rector, Thyfault et al., 2010). To investigate the effect of the SAMM50 D110G mutation on glucose metabolism, we first measured serum glucose levels in *Samm50*-KI following a 6-hour fast. Fasted *Samm50*-KI mice exhibited significantly higher serum glucose levels (Fig. 5A), although fasting insulin levels were also elevated (Fig. 5B), indicating that the increased fasting glucose was not caused by insulin deficiency. To further investigate insulin sensitivity and glucose tolerance, we conducted insulin tolerance tests (ITTs) and glucose tolerance tests (GTTs) in *Samm50*-KI mice. Compared with WT mice, serum glucose levels were significantly increased in *Samm50*-KI mice between 60 and 90 min after insulin injection and between 15 and 30 min after glucose injection (Fig. 5C-F). In conclusion, these findings indicate that mitochondrial dysfunction caused by SAMM50 D110G may lead to mild insulin resistance and glucose intolerance, possibly due to blunted cellular responses to insulin.

**Figure 5.**
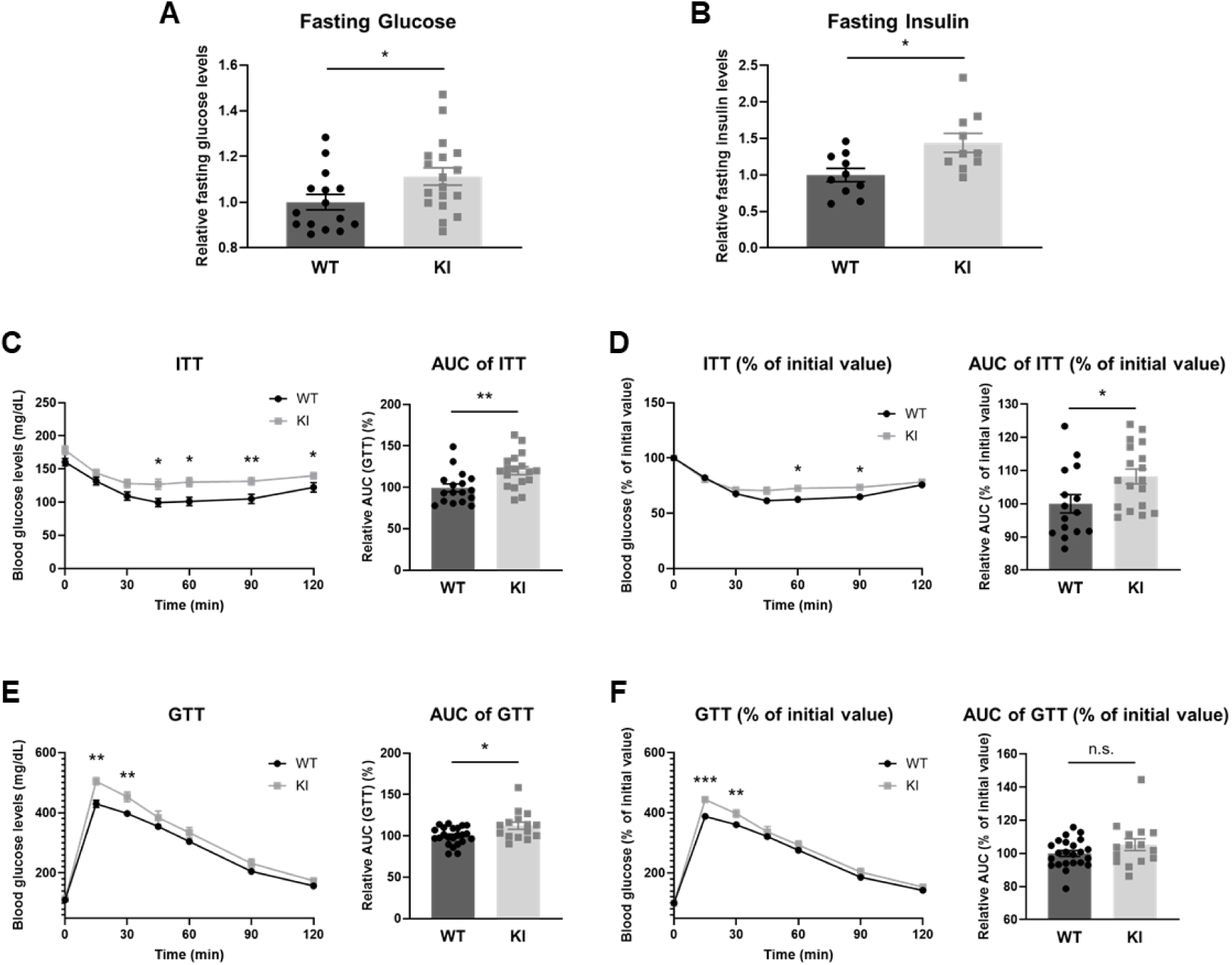
SAMM50 D110G leads to insulin resistance and glucose intolerance in mice. (**A**) Serum glucose levels (WT, n = 15; *Samm50*-KI, n = 18; *p < 0.05). (**B**) Serum insulin levels after 6 hours of fasting in WT and *Samm50*-KI mice (WT, n = 10; *Samm50*-KI, n = 10; *p < 0.05). (**C**) ITT and area under the curve (AUC) of ITT (WT, n = 15; *Samm50*-KI, n = 18; *p < 0.05; **p < 0.01). (**D**) ITT represented as % of initial glucose levels and corresponding AUC (WT, n = 15; *Samm50*-KI, n = 18; *p < 0.05). (**E**) GTT and AUC of GTT (WT, n = 23; *Samm50*-KI, n = 15; *p < 0.05; **p < 0.01). (**F**) GTT represented as % of initial glucose levels and corresponding AUC (WT, n = 23; *Samm50*-KI, n = 15; **p < 0.01; ***p < 0.001; n.s., not significant). Data are presented as means ± SEM, and statistical significance was determined using the Mann–Whitney test (**A-F**).

### A high-fat diet (HFD) triggers clinical signs of MASLD in *Samm50*-KI mice

Next, we examined whether *Samm50*-KI mice exhibited MASLD phenotypes. β-oxidation, a key pathway in hepatic lipid metabolism, is frequently impaired in MASLD (Barbier-Torres, Fortner et al., 2020). In *Samm50*-KI livers, the expression of genes involved in β-oxidation was significantly downregulated (Fig. EV3A). Although MASLD is clinically associated with weight gain and liver pathology, *Samm50*-KI mice showed no significant differences in body weight, liver weight, or liver histology (Hematoxylin and Eosin staining, H&E) compared to WT under standard diet conditions (Fig. EV3B, C). We also assessed biochemical markers that are commonly elevated in MASLD patients, including ALT, AST, triglycerides (TG), and total cholesterol (TC). Serum ALT levels were significantly increased in *Samm50*-KI mice, while AST, TG, and TC showed a trend toward elevation without reaching statistical significance (Fig. 4H; Appendix Fig. S3; Fig EV3D-E).

To directly assess the contribution of SAMM50 D110G to MASLD development, we subjected *Samm50*-KI and WT mice to an HFD. *Samm50*-KI exhibited a trend toward increased body weight during 16 weeks of HFD feeding, with significantly higher final body weight compared to WT (Fig. 6A, B; Appendix Fig. S4A-C). Liver weight and liver-to-body weight ratios were also elevated in HFD-fed *Samm50*-KI mice (Fig. 6C, D). Consistently, H&E and Oil Red O staining revealed pronounced hepatic lipid droplet accumulation in *Samm50*-KI compared with WT following HFD (Fig. 6E, F), accompanied by elevated hepatic TG levels (Fig. 6G). Serum TC and ALT levels were also increased in *Samm50*-KI mice (Fig. 6H, I). Together, these results indicate that HFD induces hepatic fat accumulation and exacerbates liver injury in *Samm50*-KI mice, leading to more severe clinical manifestations of MASLD.

**Figure 6.**
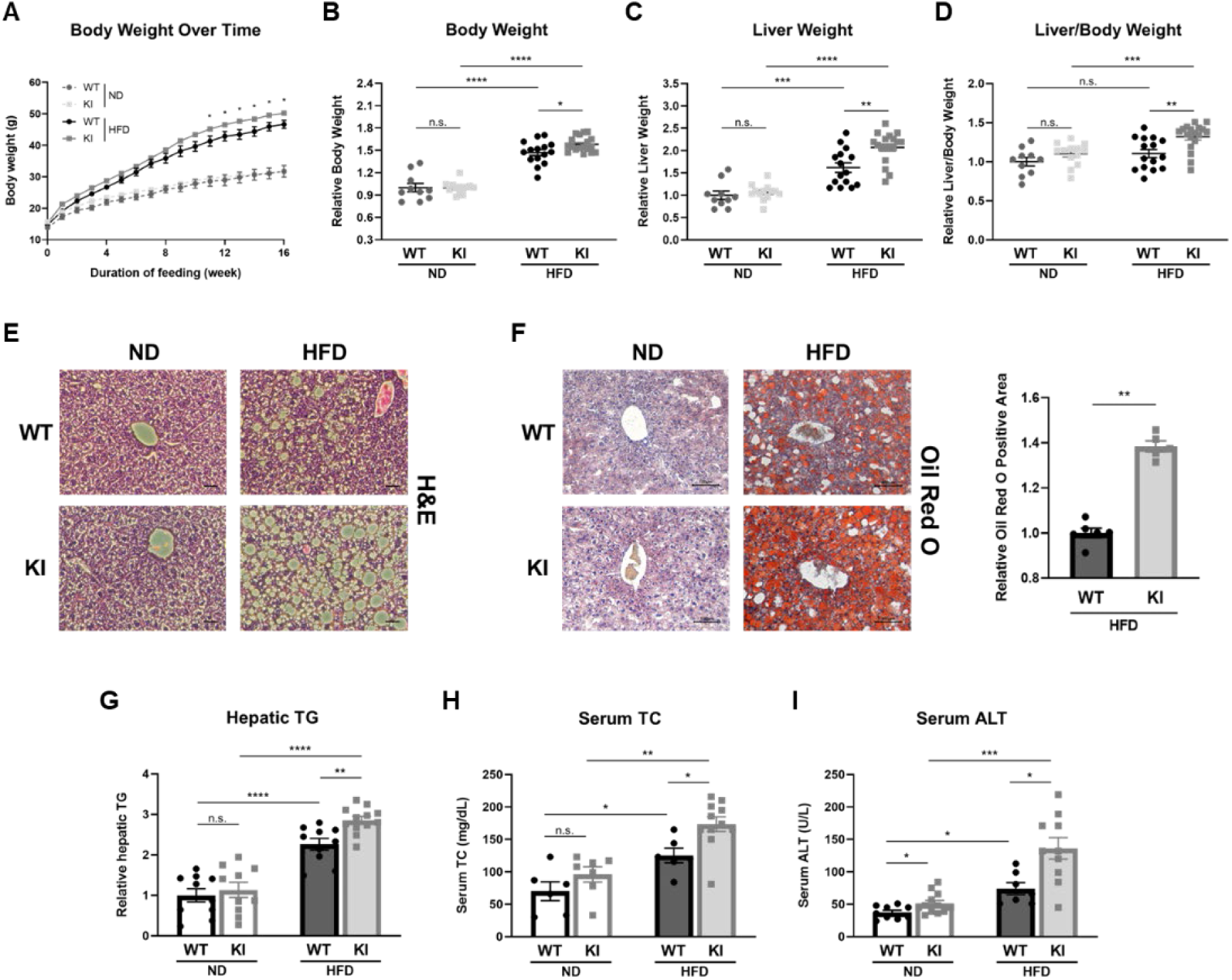
HFD triggers MASLD-associated clinical signs in *Samm50*-KI mice. **(A)** Body weight changes in WT and *Samm50*-KI mice fed either a ND or an HFD over time (WT-ND, n = 10; *Samm50*-KI-ND, n = 13; WT-HFD, n = 15; *Samm50*-KI-HFD, n = 18; *p < 0.05). (**B-D)** Comparison of body weights (**B**), liver weights (**C**), and liver-to-body weight ratios (**D**) in WT and *Samm50*-KI mice after feeding either a ND or HFD (WT-ND, n = 10; *Samm50*-KI-ND, n = 13; WT-HFD, n = 15; *Samm50*-KI-HFD, n = 18; *p < 0.05; **p < 0.01; ***p < 0.001; ****p < 0.0001; n.s., not significant). (**E)** Representative images of H&E-stained liver sections from ND- or HFD-fed mice (magnification, 200x). Scale bars: 10 μm. (**F)** Left: Representative Oil Red O-stained liver sections from WT and *Samm50*-KI mice showing lipid accumulation (magnification, 200x). Scale bars: 100 μm. Right: Quantification of Oil Red O-positive areas in livers from HFD-fed WT and *Samm50*-KI mice (WT-HFD, n = 4; *Samm50*-KI-HFD, n = 4; *p < 0.05). (**G)** Hepatic TG levels of WT and *Samm50*-KI mice (WT-ND, n = 10; *Samm50*-KI-ND, n = 10; WT-HFD, n = 10; *Samm50*-KI-HFD, n = 12; **p < 0.01; ****p < 0.0001; n.s., not significant). (**H, I)** Serum TC (**H**) (WT-ND, n = 6; *Samm50*-KI-ND, n = 7; WT-HFD, n = 6; *Samm50*-KI-HFD, n = 11; *p < 0.05; **p < 0.01; n.s., not significant), and ALT levels (**I**) of WT and *Samm50*-KI mice (WT-ND, n = 9; *Samm50*-KI-ND, n = 12; WT-HFD, n = 7; *Samm50*-KI-HFD, n = 10; (*p < 0.05; ***p < 0.001; n.s., not significant). Data are presented as means ± SEM, and statistical significance was determined using the Mann–Whitney test (**A-D**, **F-I**).

### SAMM50 D110G promotes insulin resistance, glucose intolerance, and lipid accumulation following HFD

To further assess systemic glucose homeostasis associated with MASLD progression, we performed GTT and ITT in both normal diet (ND)- and HFD-fed mice. As shown in Fig. 5, *Samm50*-KI mice exhibited mild impairments in insulin sensitivity and glucose tolerance under standard diet conditions (Fig. 5C-F). Consistently, HFD-fed WT and *Samm50*-KI mice exhibited higher serum glucose and insulin levels compared to ND-fed controls (Fig. 7A, B). Notably, these levels were significantly elevated in *Samm50*-KI mice relative to HFD-fed WT mice. Following HFD feeding, both WT and *Samm50*-KI mice displayed insulin resistance and glucose intolerance compared to their ND-fed counterparts (Fig. 7C–F). These metabolic abnormalities were more pronounced in HFD-fed *Samm50*-KI mice than in HFD-fed WT mice, indicating that the SAMM50 D110G mutation exacerbates insulin resistance and glucose intolerance under high-fat diet conditions.

**Figure 7.**
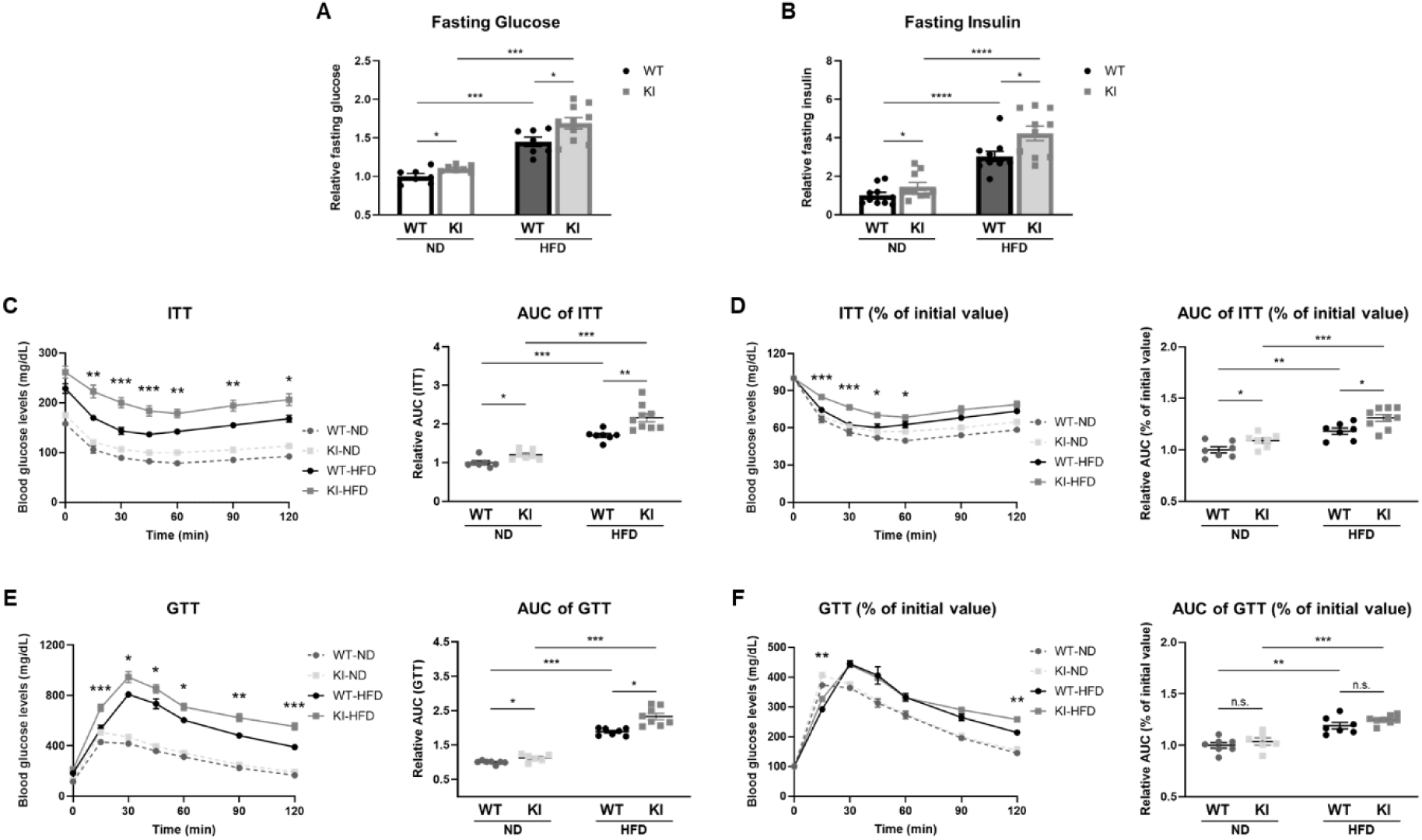
Insulin resistance and glucose intolerance in HFD-fed *Samm50*-KI mice. (**A**) Serum glucose levels (WT-ND, n = 7; *Samm50*-KI-ND, n = 7; WT-HFD, n = 7; *Samm50*-KI-HFD, n = 9; *p < 0.05; ***p < 0.001). (**B**) Serum insulin levels after 6 hours of fasting (WT-ND, n = 10; *Samm50*-KI-ND, n = 10; WT-HFD, n = 10; *Samm50*-KI-HFD, n = 10; *p < 0.05; ****p < 0.0001). **(C)** ITT and area under the curve (AUC) of ITT (WT-ND, n = 7; *Samm50*-KI-ND, n = 7; WT-HFD, n = 7; *Samm50*-KI-HFD, n = 9; *p < 0.05; **p < 0.01; ***p < 0.001). (**D)** ITT represented as % of initial glucose levels and corresponding AUC. (**E)** GTT and AUC of GTT (WT-ND, n = 7; *Samm50*-KI-ND, n = 6; WT-HFD, n = 7; *Samm50*-KI-HFD, n = 8; *p < 0.05; **p < 0.01; ***p < 0.001; n.s., not significant) (**F)** GTT represented as % of initial glucose levels and corresponding AUC. Data are presented as means ± SEM, and statistical significance was determined using the Mann–Whitney test (**A-F**).

We next investigated the molecular mechanisms underlying lipid accumulation, focusing on the involvement of mitochondria in cholesterol metabolism and lipogenesis (Ferramosca, Savy et al., 2006, Goicoechea, Conde de la Rosa et al., 2023). Genes involved in mitochondrial structure and function, including *Samm50*, *Tom40*, *Opa1*, and *Pink1*, were downregulated in HFD-fed *Samm50*-KI compared with HFD-fed WT (Fig. EV4A). In association with the increase in serum TC levels, we found an elevation in the expression of *Lss*, *Cyp51*, *Tm7sf2*, and *Dhcr24*, encoding proteins essential for cholesterol synthesis, in HFD-fed *Samm50*-KI (Fig. EV4B). Additionally, genes involved in de novo lipogenesis, including *Mlxipl*, *Srebp1*, *Acss2*, and *Fas*, were upregulated in *Samm50*-KI (Fig. EV4C), indicating that mitochondrial dysfunction in *Samm50*-KI may promote hepatic lipogenesis. Taken together, these findings suggest that aberrant mitochondrial function resulting from the SAMM50 D110G mutation, together with HFD, promotes insulin resistance and lipogenesis, thereby driving MASLD progression.

## Discussion

Several clinical studies have demonstrated an association between *SAMM50* SNPs and MASLD, and in vitro studies utilizing *SAMM50* knockdown cells have shown that SAMM50 deficiency leads to increased lipid accumulation (Li et al., 2021). However, the molecular mechanisms underlying the association between *SAMM50* rs3761472 and MASLD remain unclear. To investigate these mechanisms in vivo, we established a murine model of *SAMM50* rs3761472 variant using CRISPR/Cas9. Surprisingly, a single amino acid substitution in SAMM50 resulted in profound systemic effects in mice, affecting essential mitochondrial functions and lipid metabolism. Although it is relatively rare for a single amino acid mutation in mice to significantly alter gene expression levels (Chiu, Weng et al., 2020, Zhu, Wang et al., 2018), this mutation led to reduced SAMM50 expression in the livers of *Samm50*-KI. Considering the decreased expression of genes related to mitochondrial structure and function, D110G may alter retrograde signaling, a mitochondria-to-nucleus communication pathway through which mitochondrial dysfunction modulates nuclear gene transcription (Butow & Avadhani, 2004, Warnsmann, Marschall et al., 2022). This dysregulation could reflect structural abnormalities in mitochondria, which may be further examined in future studies using transmission electron microscopy.

Interestingly, *SAMM50* knockdown enhances mitophagy in vitro (Jian, Chen et al., 2018), whereas *Samm50*-KI mice exhibited reduced mitophagy, suggesting distinct molecular consequences of the SAMM50 alteration. While *SAMM50* knockdown inhibits PINK1 processing and degradation, leading to its accumulation (Jian et al., 2018), the D110G mutation may induce structural alterations in the SAM complex that impair OMM complex assembly or mitochondrial protein import, thereby preventing proper PINK1 stabilization on the OMM. Consistent with our observations, impaired mitophagy and decreased PINK1/Parkin expression have also been reported in the MASLD model (Yao, Li et al., 2023).

The SAMM50 D110G mutation compromises ATP generation and promotes ROS production, triggering immune responses, ultimately leading to hepatocellular death, insulin resistance, and glucose intolerance. Moreover, hepatic steatosis, hepatic TG, serum TC, and ALT levels were significantly increased in HFD-fed *Samm50*-KI. Given the role of mitochondrial dysfunction in MASLD, we examined sex-specific effects of the SAMM50 D110G, as females are generally more resistant to MASLD due to sex hormone-mediated protection and enhanced mitochondrial function (McCoin, Von Schulze et al., 2019, Yuan, Kardashian et al., 2019). In contrast to *Samm50*-KI males, females showed no significant alterations in mitochondrial gene expression or ALT levels, suggesting protection against mitochondrial dysfunction and liver injury (Appendix Fig. S5A-H). Consequently, the SAMM50 D110G is associated with an increased risk of MASLD, particularly in males. These mechanisms underscore the importance of mitochondrial health in MASLD development and progression and offer valuable insights into the pathogenesis of the disease.

Although this study focused on the liver, previous studies have shown that mitochondrial gene deficiencies impair function in other tissues, particularly the brain and heart (Shirakabe, Zhai et al., 2016, Song et al., 2017, Torres-Odio, Key et al., 2017). To explore potential extrahepatic effects of the SAMM50 D110G variant, we assessed *Samm50* expression in the brain, heart, kidney, and spleen. While a 25% reduction was observed in the brain, no significant changes were detected in other tissues (Fig. EV5A-D). These findings support the liver as the primary target organ, although potential effects in the brain require further investigation.

Our study demonstrates that the *SAMM50* SNP rs3761472 impairs mitochondrial function, thereby contributing to the development of MASLD. Using CRISPR/Cas9 genome editing, we generated *Samm50*-KI mice harboring the rs3761472 variant, which displayed mitochondrial dysfunction and key pathological features of MASLD (Fig. 8). These findings provide new mechanistic insights into the pathogenic role of rs3761472 and underscore its potential as a target for therapeutic intervention and precision medicine. Importantly, rs3761472 is also associated with increased risks of cirrhosis and hepatocellular carcinoma, suggesting broader relevance in chronic liver diseases. Targeting this variant may enhance mitochondrial function or serve as a biomarker for identifying high-risk individuals and tailoring personalized treatment strategies.

**Figure 8.**
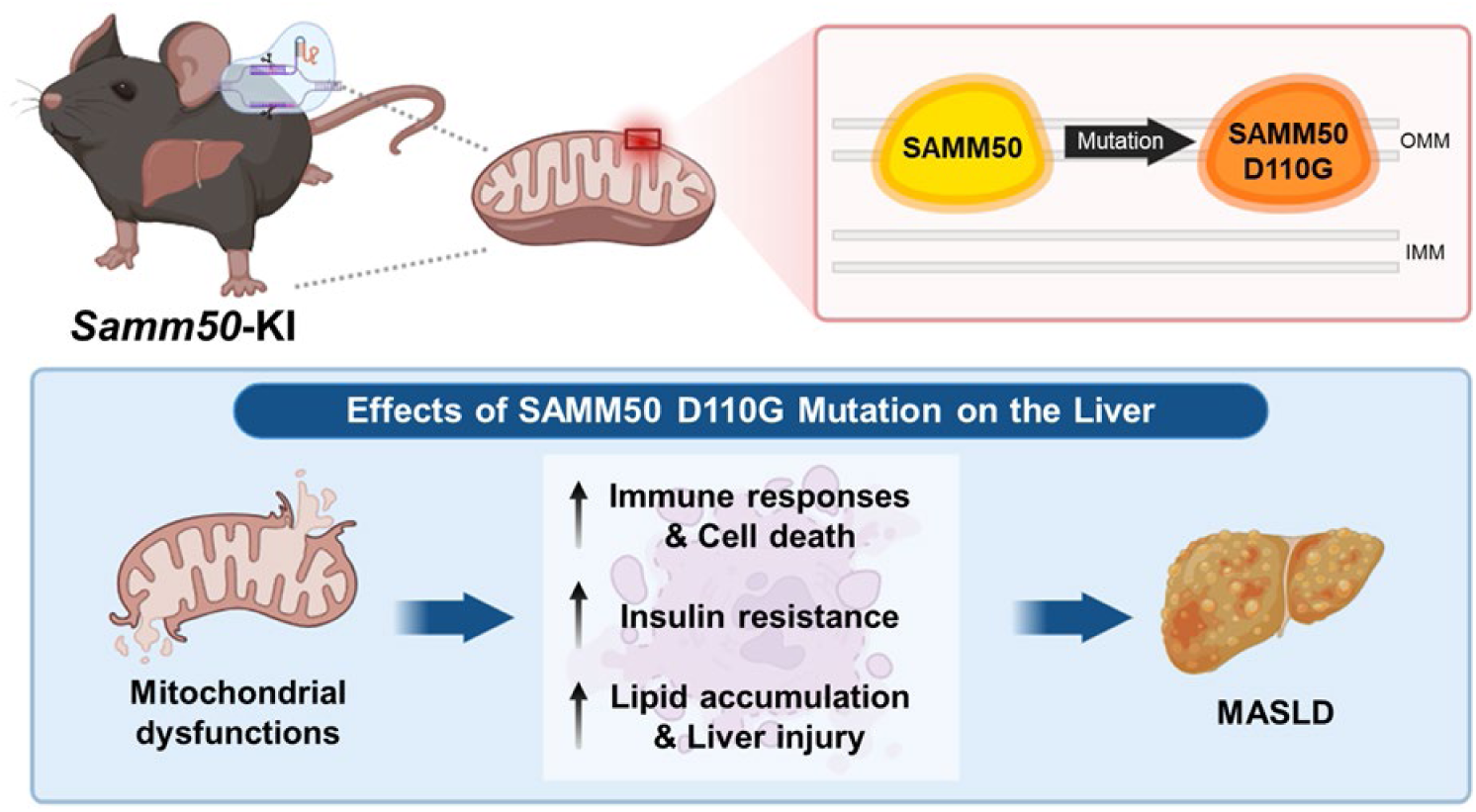
Working model of SAMM50 D110G functions in the liver. CRISPR/Cas9–generated *Samm50*-KI mice carrying the rs3761472 variant exhibit mitochondrial dysfunction and key pathological features of MASLD. The model illustrates how the SAMM50 mutation impairs mitochondrial quality control, contributing to hepatic lipid accumulation and liver injury.

## Methods

### Genetic association analysis

Previously, a GWAS of ALT levels was conducted using 6,949 individuals genotyped using the Korea Biobank Array optimized for the Korean population (Moon, Kim et al., 2019). Among the associations found, a nonsynonymous SNP, rs3761472, in *SAMM50* was significantly associated with ALT levels (β = 0.0362, P = 7.23 × 10^-6^). Based on the annotation from dbNSFP v2.7, in silico prediction algorithms predicted that SNP rs3761472 would be damaging (see Appendix Table S1). In silico prediction tools from dbNSFP v2.7 categorized rs3761472 as damaging, with predictions of “Damaging,” “Disease-causing,” or a CADD score > 15 suggesting functional impairment of *SAMM50* (Appendix Table S1). We further analyzed genotype dosages for rs3761472 in 151,912 Korean-ancestry individuals from the Korea Biobank Array to assess associations with serum liver enzymes including ALT, AST, and γ-glutamyl transferase (GGT), as well as lipid measures including high-density lipoprotein cholesterol (HDL-C), low-density lipoprotein cholesterol (LDL-C), TG, and TC (Kim, Moon et al., 2022). Among the traits, ALT, AST and TG were natural-log transformed to approximate normality. Linear regression under an additive genetic model was performed in R (v4.2.0) adjusting for age, age², sex, body-mass index (BMI), recruitment region, and the first four genetic principal components. Finally, we queried the Biobank Japan PheWeb for associations of rs3761472 with liver related disease phenotypes (Sakaue et al., 2021).

### Mice

Male WT C57BL/6 mice were purchased from Orient Bio (Seongnam, South Korea). First-generation (F1) *Samm50*-KI mice were generated by GEMCRO (formerly Toolgen; Seoul, Korea) using the CRISPR/Cas9 system from a C57BL/6 founder obtained from Orient Bio. To minimize environmental variability, WT and *Samm50*-KI mice were housed under identical environmental conditions throughout the experimental period. Mice were housed in groups of up to five animals per cage in a specific pathogen-free environment, maintained under a 12-hour light/dark cycle at room temperature (20–25°C) and 30–70% relative humidity, with ad libitum access to food and water. For the experiments, 16-week-old WT and homozygous KI mice were sacrificed, and serum and liver samples were collected. Mice were bred in the animal care facility at Konkuk University (Seoul, Korea) and used in accordance with a protocol approved by the Institutional Animal Care and Use Committee (IACUC).

### Generation of a CRISPR/Cas9–mediated KI mouse line and genotyping

F1 mice with identical genotypes were interbred to produce F2 homozygous mice. Genotyping was performed by polymerase chain reaction (PCR) amplification of genomic DNA isolated from the tail tip, followed by Sanger sequencing. The details of the PCR and sequencing primers are provided in Appendix Table S2.

### Quantitative reverse transcription-PCR (RT-qPCR)

Total RNA was isolated from the liver tissues of WT or *Samm50*-KI mice using the PureLink RNA Mini Kit (Invitrogen, Carlsbad, CA, USA) according to the manufacturer’s instructions. The quantity and quality of isolated RNA were analyzed using a NanoPhotometer N60/N50 spectrophotometer (Implen, München, Germany). cDNA was synthesized from total RNA using the SuperScript III First-Strand Synthesis Supermix (Invitrogen). qPCR was performed on a LightCycler 96 system (Roche, Mannheim, Germany) using SsoAdvanced Universal SYBR Green Supermix (Bio-Rad, Hercules, CA, USA). Gene-specific primers used for qPCR are listed in Appendix Table S3. Gene expression levels were normalized to those of glyceraldehyde-3-phosphate dehydrogenase (*Gapdh*).

### Western blotting

Western blotting was performed using two distinct protocols. For MTX1, PINK1, and VDAC, liver lysates were prepared with NP-40 extraction buffer, and protein concentrations were determined using a BSA assay. Samples and Precision Plus Protein Kaleidoscope Standards (Bio-Rad) were resolved on 4–20% Mini-PROTEAN TGX gels (Bio-Rad) and transferred to PVDF membranes. Membranes were incubated overnight with primary antibodies, washed with TBST, and probed with anti-mouse or anti-rabbit secondary antibodies (Cell Signaling Technology, Danvers, MA, USA) for 1–2 hours. Signals were detected using the Fusion FX system (Viber, Loumat, France). For all other proteins, liver lysates were extracted with RIPA buffer containing protease inhibitors (Roche, Basel, Switzerland), and proteins were separated on 10% or 15% SDS-PAGE gels with PageRuler™ Plus markers (Thermo Fisher Scientific, Waltham, MA, USA), transferred to nitrocellulose membranes, and detected via enhanced chemiluminescence (Thermo Fisher Scientific) using the ChemiDoc system (Bio-Rad). Antibodies used are listed in Appendix Table S4.

### In silico protein structure prediction

Protein structure predictions for SAMM50 and its D110G mutant were performed using ColabFold v1.5.5, a Google Colab-based implementation of AlphaFold2. Structural alignment and residue labeling were performed using ChimeraX (version 1.9). Pairwise alignment using the MatchMaker tool quantified structural differences using the root-mean-square deviation (RMSD). The positions of the D110 residue and its mutated counterpart (G110) were identified and visualized using 3D structures.

### Measurement of NAD levels and NAD^+^/NADH ratio

NAD^+^ and NADH levels and NAD^+^/NADH ratios were measured using an NAD/NADH Assay Kit (#ab65348; Abcam, Cambridge, MA, USA) as described by the manufacturer. Briefly, tissue samples were homogenized in NADH/NAD extraction buffer and filtered through a 10-kD spin column (#ab93349; Abcam) to remove the enzymes. After completing the assay, the absorbance was measured at 450 nm using a microplate reader (Epoch; BioTek, Winooski, VT, USA). The final protein concentrations were normalized to the liver protein concentration, as determined by the Bradford assay.

### ATP assay

ATP levels in mouse liver were measured using a colorimetric ATP assay kit (#ab83355; Abcam) according to the manufacturer’s instructions. Liver tissue lysates were deproteinized with perchloric acid and neutralized with potassium hydroxide. Absorbance was measured at 570 nm using a microplate reader (Epoch; BioTek) to quantify total ATP levels.

### Measurement of mtROS

mtROS production in liver tissues was measured using MitoSOX Mitochondrial Superoxide Indicators (#M36008; Invitrogen). Paraffin-embedded liver tissue (4 μm thick) and frozen liver tissue (5 μm thick) sections were incubated with MitoSOX Red (5 μM) in the dark at 37°C for 15 min. The sections were then stained with ProLong Gold Antifade Mountant with DAPI (Invitrogen) in the dark at 25°C for 1 min. The sections were imaged using a fluorescence microscope (Nikon Eclipse Ts2R; Nikon, Tokyo, Japan) and the fluorescence intensity was quantified using ImageJ software (Media Cybernetics, Rockville, MD, USA). Three fields of view were imaged for each sample.

### Immunofluorescence

Formalin-fixed paraffin-embedded mouse liver sections (4 µm) were deparaffinized with xylene and ethanol. Antigen retrieval was performed using a TintoRetriever Heat Retrieval System (Bio SB, USA) in 10 mM sodium citrate buffer (pH 6.0) for 10 minutes. The sections were blocked at 25°C for 1 h and incubated overnight at 4°C with an anti-Nos2 antibody (#sc-7271; Santa Cruz Biotechnology, USA). After washing with PBS, tissues were incubated with Alexa Fluor 488-conjugated anti-mouse IgG (#R37120; Invitrogen) for 1 h at 25°C. Sections were mounted using ProLong Gold Antifade Mountant with DAPI (Invitrogen) and imaged using a fluorescence microscope (Nikon Eclipse Ts2R; Nikon). The fluorescence intensity was quantified using ImageJ software. Three fields per sample were analyzed.

### TUNEL assay

DNA fragmentation in apoptotic cells within liver sections was visualized by terminal deoxynucleotidyl transferase-mediated dUTP nick-end labeling (TUNEL) using an in situ BrdU Red DNA Fragmentation Assay Kit (#ab66110; Abcam) according to the manufacturer’s instructions. The slides were counterstained with ProLong Gold Antifade Mountant with DAPI (Invitrogen) in the dark at 25°C for 1 min. TUNEL-positive red fluorescence signals were analyzed using a fluorescence microscope (Nikon Eclipse Ts2R; Nikon). Three fields per sample were imaged for quantification of TUNEL-positive cells.

### ITTs and GTTs

ITTs were conducted on mice that had fasted for 6 h; GTTs were performed on mice that had fasted for 16 h. Baseline blood glucose levels in the tail vein blood samples were measured using a glucometer. For ITTs, human insulin was administered via intraperitoneal injection at a dose of 0.75 IU/kg body weight, and a 20% glucose solution was administered in the same manner at a dose of 2 g/kg body weight. Glucose levels were measured 15, 30, 45, 60, 90, and 120 min after insulin injection for ITTs and after glucose injection for GTTs.

### Measurement of serum insulin levels

Serum insulin levels were determined using a rat/mouse insulin ELISA kit (#EZRMI-13K; Millipore, MA, USA) according to the manufacturer’s instructions.

### Biochemical analysis

Mouse serum was obtained by centrifuging whole blood samples at 2,000 × g for 20 min, and collecting the supernatant. Serum levels of ALT, AST, TC, and TG were measured using an automated dry chemistry analyzer (FUJI DRI-CHEM 7000i, Fujifilm, Tokyo, Japan).

### HFD

C57BL/6 and *Samm50*-KI male mice, aged four weeks at the start of the study, were used for the experiments. Both WT and *Samm50*-KI mice were maintained on a ND (D12450B; Research Diets, New Jersey, USA) or HFD containing 60% kilocalories of fat (D12492; Research Diets, New Jersey, USA) for 16 weeks. Body weight was monitored weekly throughout the experiment.

### Histopathological analysis

The liver tissues were fixed in 10% formalin, dehydrated, and embedded in paraffin. Tissues were sectioned at a thickness of 4 µm for H&E and Oil Red O staining, which were performed by Labcore (Labcore, Seoul, Korea). Each group included four independent replicates, with at least two fields analyzed per replicate.

### Measurement of Hepatic TG

Hepatic TG content was quantified using the PicoSens Triglyceride Assay Kit (#BM-TGR-100; BIOMAX, Seoul, Korea), according to the manufacturer’s instructions.

### Statistical analyses

Data are presented as mean ± standard error of the mean (SEM) of independent biological replicates. All statistical analyses were performed using the Mann–Whitney test.

## Acknowledgements

We thank Dr. J.M.K. from the National Institute of Health of Korea for providing constructive advice during the early stage of this project.

## Author contributions

**Suyeon Kim**: Conceptualization; Data curation; Formal analysis; Methodology; Project administration; Validation; Visualization; Writing – original draft. **Nahyun Kim**: Investigation; Validation. **Uijin Kim**: Investigation; Validation. **Young Jin Kim**: Formal analysis; Methodology; Software. **Jun Ho Yun**: Investigation; Validation. **Jiwon Heo**: Investigation.

**Hyunwoo Lee**: Investigation. **Jiwon Choi**: Investigation. **Inhae Jeong**: Investigation. **Bong-Jo Kim**: Conceptualization; Funding acquisition; Project administration; Resources; Supervision. **Ha Youn Shin**: Conceptualization; Funding acquisition; Project administration; Resources; Supervision; Writing – review & editing.

## Data Availability

This study includes no data deposited in external repositories.

---

**Figure EV1.**
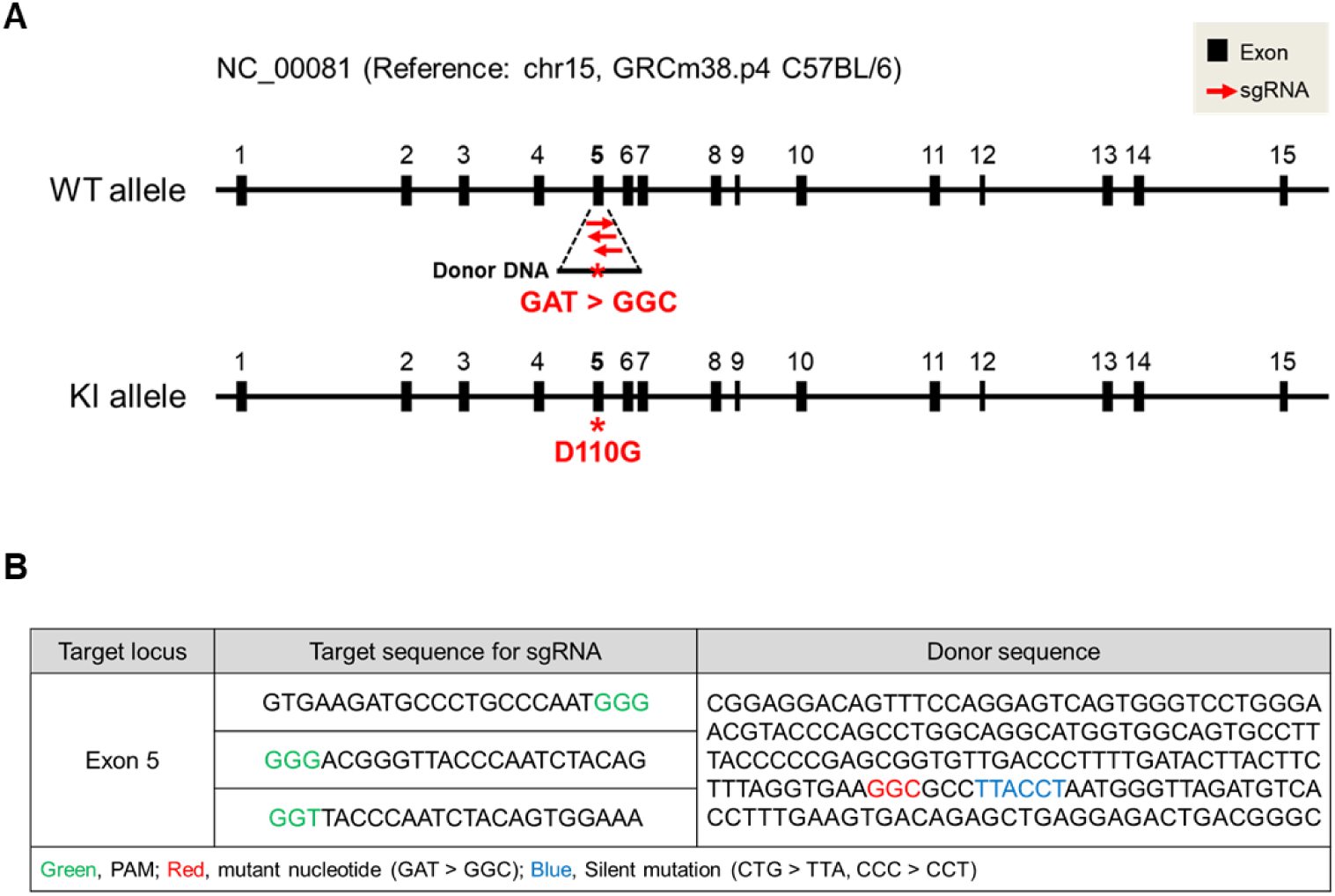
Experimental designs for generating SAMM50 D110G mutant mice using the CRISPR/Cas9 system. (A) Genomic locus of the *Samm50* gene in mice, showing the binding sites of gRNAs on the target region. In the knock-in mice, the aspartic acid at position 110 (D110) encoded by exon 5 of *Samm50* is converted to glycine (G). (B) sgRNAs and donor sequences used to introduce the SAMM50 D110G mutation. The protospacer adjacent motif (PAM) is highlighted in green, target nucleotide changes are shown in red, and silent mutations to prevent off-target Cas9 cutting are marked in blue. To introduce the D110G mutation, we modified exon 5 of *Samm50* by substituting the GGC with the original GAT codon. We also introduced silent mutations, CTG to TTA and CCC to CCT, in the adjacent target sequence to eliminate non-target PAM sequences, reducing the likelihood of off-target cleavage by Cas9 nuclease.

**Figure EV2.**
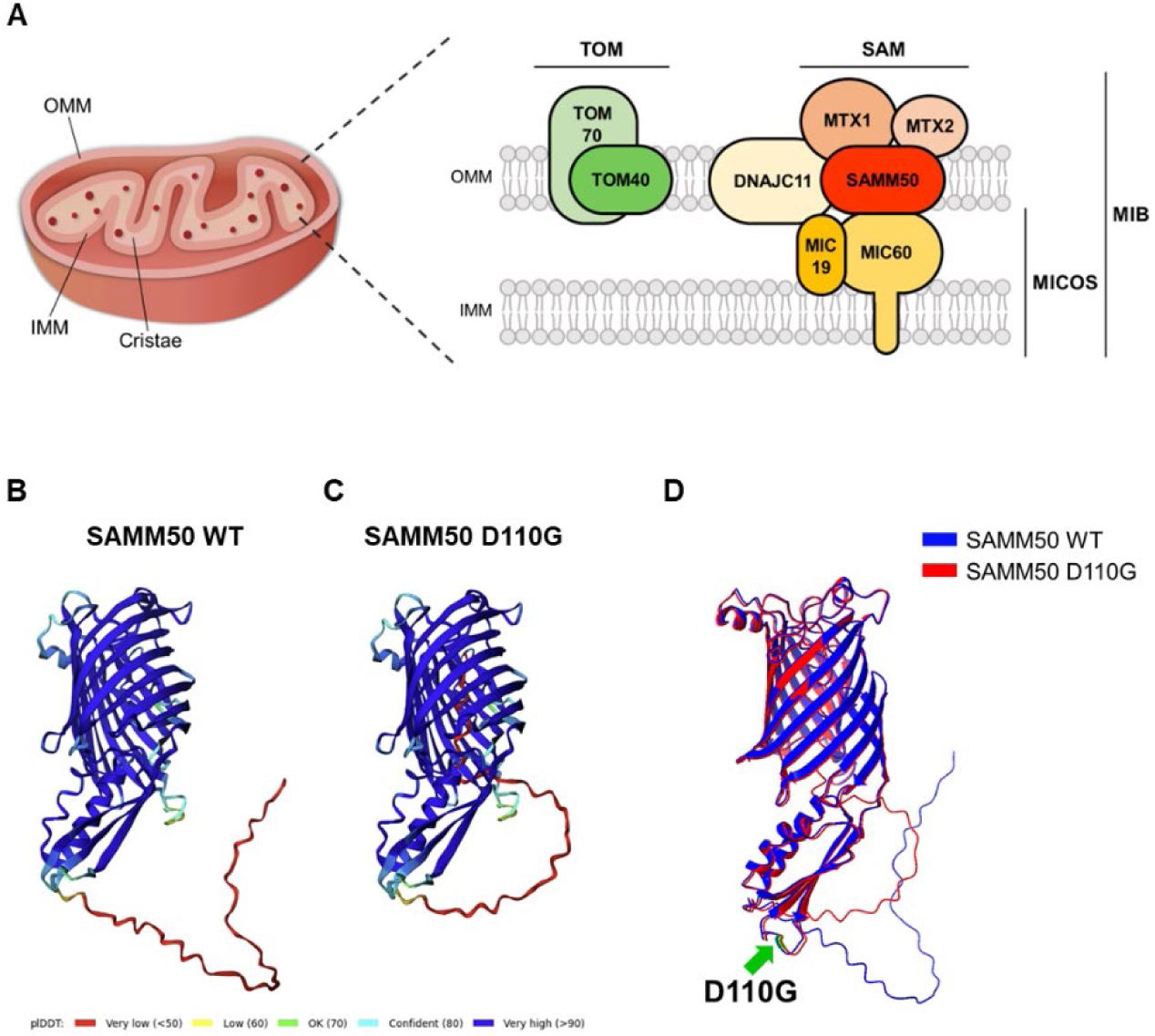
Essential mitochondrial structural components and predicted protein structures of SAMM50 WT and D110G mutant. (**A**) Schematic illustration of mitochondrial structures and key mitochondrial structural protein components. Mitochondria are enclosed by a double membrane structure. The outer mitochondrial membrane (OMM) forms the outermost layer, whereas the inner mitochondrial membrane (IMM) folds into numerous cristae. Mitochondrial integrity is maintained by essential protein complexes. The translocase of the outer membrane (TOM) and sorting and assembly machinery (SAM) complexes are located in the OMM, whereas the mitochondrial contact site and cristae organizing system (MICOS) is in the IMM; mitochondrial intermembrane space bridging (MIB) complexes mediate interactions between the OMM and IMM. The TOM complex includes TOM40 and TOM70, whereas the SAM complex contains SAMM50, MTX1, and MTX2. The MICOS complex consists of MIC19 and MIC60, and the MIB complex includes DNAJC11, SAM, and MICOS components. MIB complexes are crucial for maintaining cristae structure. (**B, C**) Predicted protein structures of SAMM50 WT (**B**) and SAMM50 D110G mutant (**C**) generated using ColabFold v1.5.5, which implements AlphaFold2. (**D**) Structural alignment of the two protein models displays the SAMM50 WT in blue and the D110G mutant in red. Residues D110 (WT) and G110 (mutant) are indicated with the green arrow for clarity.

**Figure EV3.**
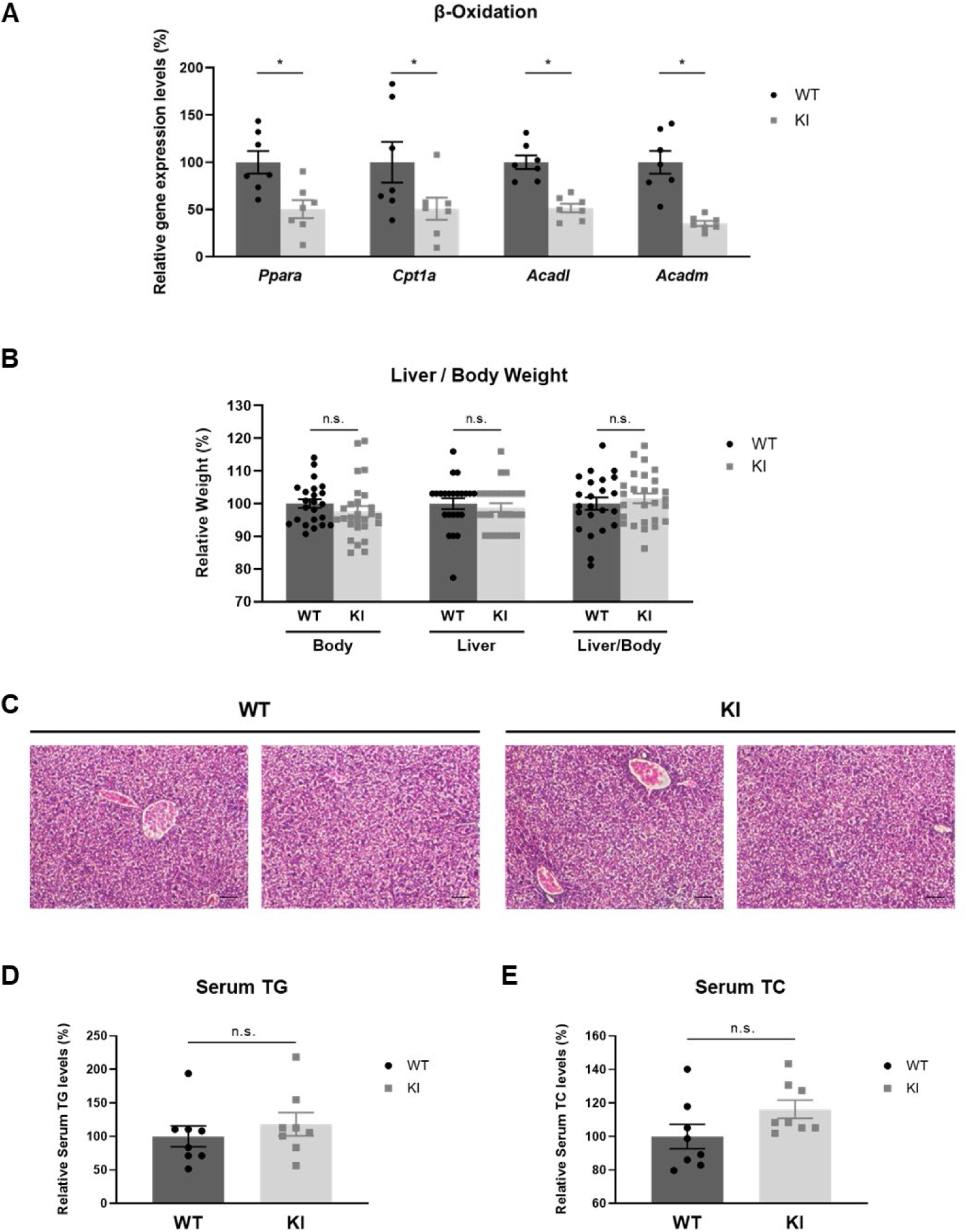
Examination of MASLD features in *Samm50*-KI mice. (**A**) Relative mRNA expression levels of β-oxidation–related genes (*Ppara*, *Cpt1a*, *Acadl*, and *Acadm*) in liver tissues from WT and *Samm50*-KI mice (WT, n = 7; *Samm50*-KI, n = 7; *p < 0.05). Gene expression was normalized to *Gapdh* and is presented relative to WT levels. (**B**) Comparison of body weights, liver weights, and liver-to-body weight ratios between WT and *Samm50*-KI mice (WT, n = 23; *Samm50*-KI, n = 27; n.s., not significant). (**C**) Representative images of H&E-stained liver sections from WT and *Samm50*-KI mice (magnification, 200x). Scale bars: 10 μm. (**D, E**) Relative levels of serum triglycerides (TG) (**D**), and total cholesterol (TC) in WT and *Samm50*-KI mice (**E**) (WT, n = 8; *Samm50*-KI, n = 8; n.s., not significant). Data are presented as means ± SEM, and statistical significance was determined using the Mann– Whitney test (**A**, **B**, **D**, **E**).

**Figure EV4.**
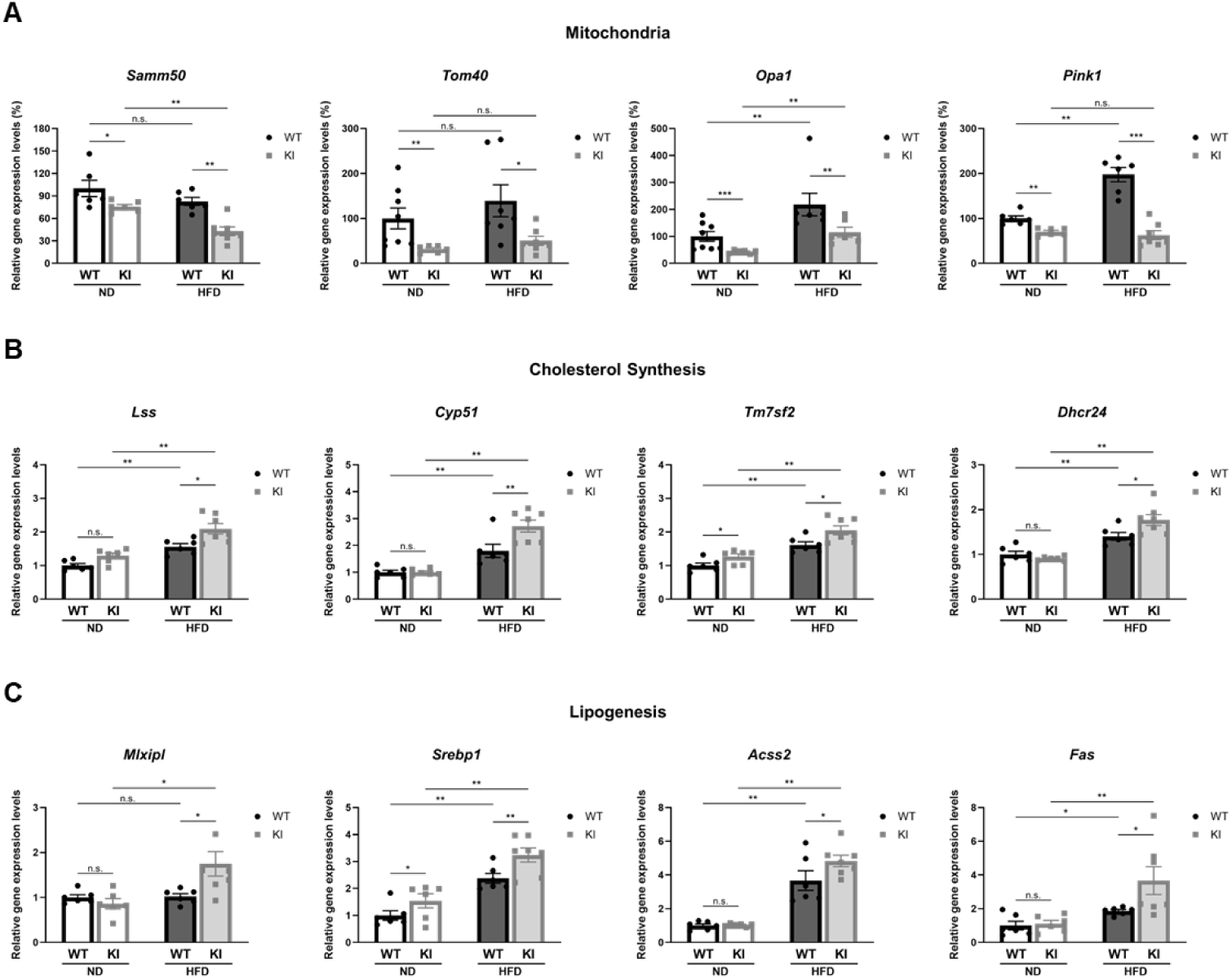
Downregulation of mitochondria-associated genes promotes cholesterol synthesis and liver lipogenesis in HFD-fed *Samm50*-KI mice. (**A**) Relative expression levels of the mitochondrial structure- and function-related genes, *Samm50*, *Tom40*, *Opa1*, and *Pink1* (WT-ND, n ≥ 6; *Samm50*-KI-ND, n ≥ 6; WT-HFD, n ≥ 6; *Samm50*-KI-HFD, n ≥ 6; *p < 0.05; **p < 0.01; ***p < 0.001; n.s., not significant). (**B**) Relative expression levels of the cholesterol synthesis pathway-related genes, *Lss*, *Cyp51*, *Tm2sf7*, and *Dhcr24* (WT-ND, n ≥ 6; *Samm50*-KI-ND, n ≥ 6; WT-HFD, n ≥ 6; *Samm50*-KI-HFD, n ≥ 6; *p < 0.05; **p < 0.01; n.s., not significant). (**C**) Relative expression levels of the de novo lipogenesis-related genes, *Mlxipl*, *Srebp1*, *Acss2*, and *Fas*. (WT-ND, n ≥ 6; *Samm50*-KI-ND, n ≥ 6; WT-HFD, n ≥ 6; *Samm50*-KI-HFD, n ≥ 6; *p < 0.05; **p < 0.01; n.s., not significant). mRNA levels of each gene were normalized to that of *Gapdh* and are presented relative to mRNA levels in ND-fed WT mice. Data are presented as means ± SEM, and statistical significance was determined using the Mann–Whitney test (**A-C**).

**Figure EV5.**
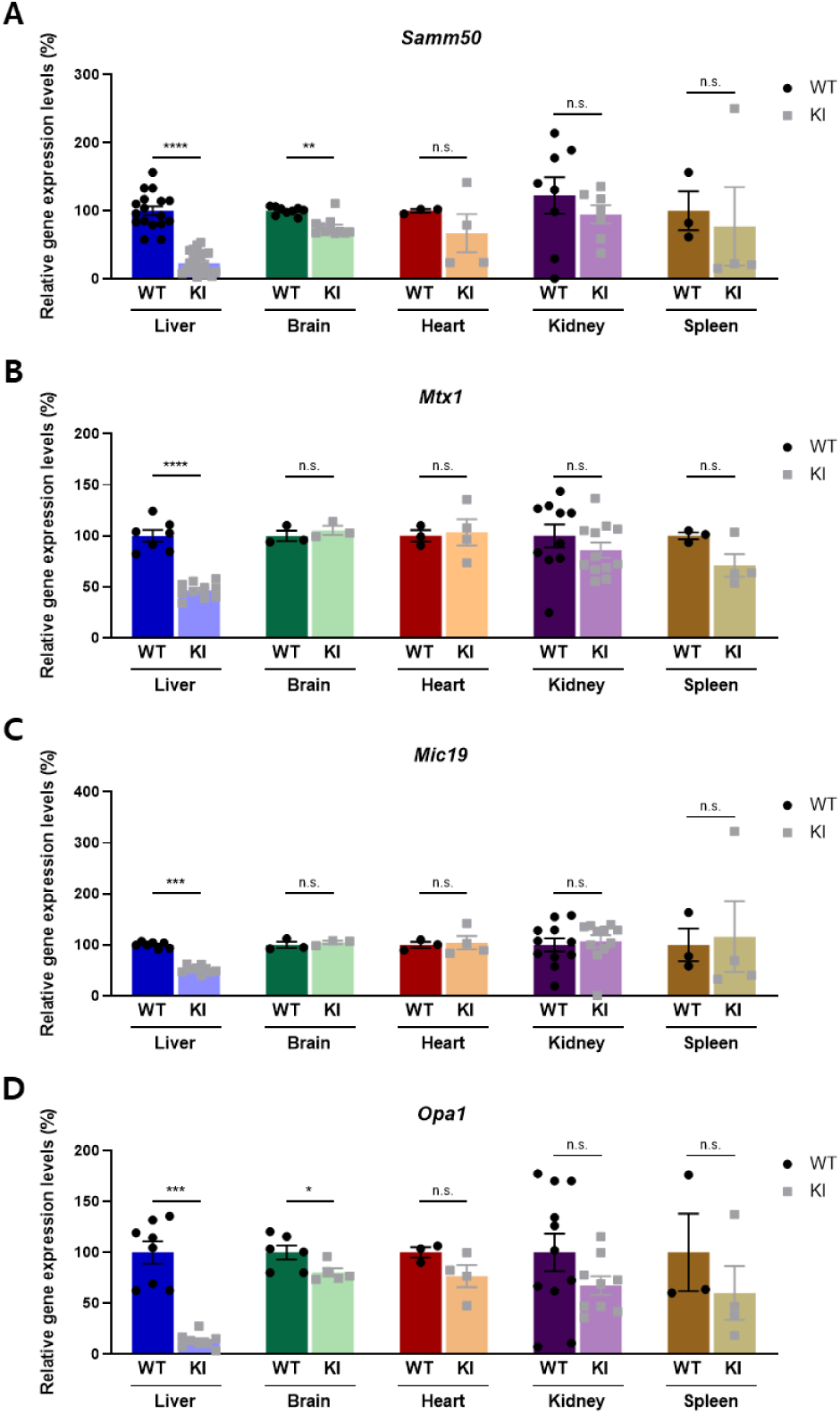
Comparative expression of mitochondrial genes across multiple tissues in WT and *Samm50*-KI mice. (**A-D**) Relative mRNA expression levels of mitochondrial-related genes: *Samm50* (**A**), *Mtx1* (**B**), *Mic19* (**C**), and *Opa1* (**D**) in the liver, brain, heart, kidney, and spleen of WT and *Samm50*-KI mice (WT, n ≥ 3; *Samm50*-KI, n ≥ 3; *p < 0.05; **p < 0.01; ***p < 0.001; n.s., not significant). mRNA levels of each gene were normalized to those of *Gapdh*, and KI mRNA levels are presented relative to those of WT. Data are presented as means ± SEM, and statistical significance was determined using the Mann–Whitney test (**A-D**).

